# How Sampling-Based Overdispersion Undermines India’s Tiger Monitoring Orthodoxy

**DOI:** 10.1101/708628

**Authors:** Arjun M. Gopalaswamy, K. Ullas Karanth, Mohan Delampady, Nils Chr. Stenseth

**Affiliations:** Statistics and Mathematics Unit, Indian Statistical Institute, Bangalore Centre, Bengaluru – 560059, India; Centre for Wildlife Studies, 37/5, Yellappa Garden, Yellappa Chetty Layout, Sivanchetti Gardens, Bengaluru, Karnataka 560042, India; Wildlife Conservation Society, Global Conservation Program, Bronx, New York 10460, USA; National Centre for Biological Sciences, Tata Institute of Fundamental Research, Bengaluru – 560065, India; Centre for Ecological and Evolutionary Synthesis (CEES), Department of Biosciences, University of Oslo, Blindernveien 31, NO-0316, Oslo, Norway

## Abstract

Conservation agencies entrusted with recovery of iconic mammals may exaggerate population trends without adequate scientific evidence. Recently, such populations were termed as ‘political populations’ in the conservation literature. We surmise that political populations emerge when agencies are pressured to report abundances at large spatial scales for species that are difficult to survey. Indian tiger conservation agencies use an experimental approach called double-sampling using index-calibration models. A recent, mathematical, study demonstrated the unreliability of this approach in the context of India’s tigers. Yet, this approach continues to be applied and even promoted by global tiger conservation agencies in other tiger range countries. In this article, we aim to: (1) discuss the ecological oddities emerging from results of India’s national tiger surveys, (2) demystify the mathematics underlying the problems of this survey methodology and (3) confront these findings with results from India’s recent national tiger survey of 2014. Our analyses show that the predictions of tiger abundance using sign-based indices reported in the 2014 survey in fact vary greatly and can be severely misleading and confirming the presence of high sampling-based overdispersion and parameter covariance. We call for species conservation initiatives to implement monitoring methods that are designed to clearly answer, a priori, scientific or management objectives instead of potentially implementing them as reactions to extraneous, social or fund raising pressures.

## INTRODUCTION AND BACKGROUND

Krebs (1991) recognized that monitoring programs must advance our knowledge of the underlying dynamics of animal populations if they are to improve either science or conservation. Towards this end, Nichols and Williams (2006) recommend *a priori* designing of animal monitoring programs to answer clearly defined scientific or management questions. And in practice, Williams et al. (2002) identify two major sources of uncertainty (imperfect detection and inappropriate spatial sampling), which must be addressed while implementing monitoring programs to generate strong inferences about animal population dynamics.

Monitoring programs for some of the world’s most iconic endangered mammals, however, appear to routinely ignore these profound insights leading to claims about population dynamics of such species resting on weak inferences and untested leap of faith arguments. For example, Darimont et al. (2018) explain how population trends reported by agencies for several charismatic carnivores lack adequate scientific support. Using case studies of wolves (*Canis lupus*) in USA and Sweden and brown bears *(Ursus arctos)* in Romania and Canada, they demonstrate how population increases claimed by federal agencies are exaggerated. Hypothesizing that these claims serve political interests, they coined the term “political populations” (Darimont et al. 2018) to identify such species. Therefore, it is important to assess whether conservation agency claims about population trends of political populations arise from poorly framed monitoring questions, inadequate sampling designs or from extraneous social considerations such as ‘motivated reasoning’ (Kunda 1990). In this essay, we attempt to disentangle these factors based on official reports of monitoring wild tigers (*Panthera tigris*) in India, also suspected to be a political population by Darimont et al. (2018).

The tiger is an ideal ‘political species’ for such an investigation because of the global attention and massive conservation investments it has attracted (PTI 2016). In this article, we first contrast official results of Indian tiger surveys with ecological theory and prior scientific knowledge of tiger population dynamics. Thereafter, we examine the underlying statistical factors leading to the scientific inferences from these surveys. Finally, we broaden implications of our results to political populations of other charismatic species.

## INDIA’S CLAIMS OF RISING TIGER NUMBERS

The Indian country-wide official surveys of 2006, 2010 and 2014 (herein referred as “NTE surveys”) report estimates of tiger population sizes at 1411 (1165-1657), 1706 (1507-1896) and 2226 (1945-2491), respectively (Jhala et al. 2011a, Jhala et al. 2011b, Jhala et al. 2015). The numbers in brackets putatively represent the range, without a clear statistical explanation of how these values are derived. Regardless, these numbers with their reported error bounds indicate significant increases in tiger numbers in India.

Considering only areas that were actually surveyed (summarized from Jhala et al. 2015), these numbers translate to a 17.2% increase in tiger abundance and a corresponding increase of 34.4% in local tiger density, implying that local tiger density *D* rose at twice the rate of tiger abundance *N* (ΔD/ΔN = 2) between 2006-2010. Between 2010-2014, an even steeper 30% increase in tiger abundance was reported, but this time there was corresponding decrease in local tiger density down to 19%. These results imply a *reversal* of the tiger population growth mechanism (from ΔD/ΔN = 2 to ΔD/ΔN = 0.63) after four years, with the year 2010 as the point of inflexion.

Furthermore, between 2006-2010, the surveys reported a simultaneous contraction of tiger range by 12.9% (or 11,400 km^2^). In contrast the results of the next survey interval (2010-2014) claim an abrupt *reversal* of the earlier pattern, reporting a range expansion of 9.4%, suggesting a tiger re-colonization of 7,250 km^2^ of new habitat (computed from Jhala et al. 2015).

These tiger population increase mechanisms imply a concave *upward* relationship between tiger abundance and occupancy (Figure 1) and stand in contrast to the general mechanism of a monotonically increasing, but concaving *downward*, relationship based on basic scientific literature (see Gaston et al. 2000).

**Figure 1.**
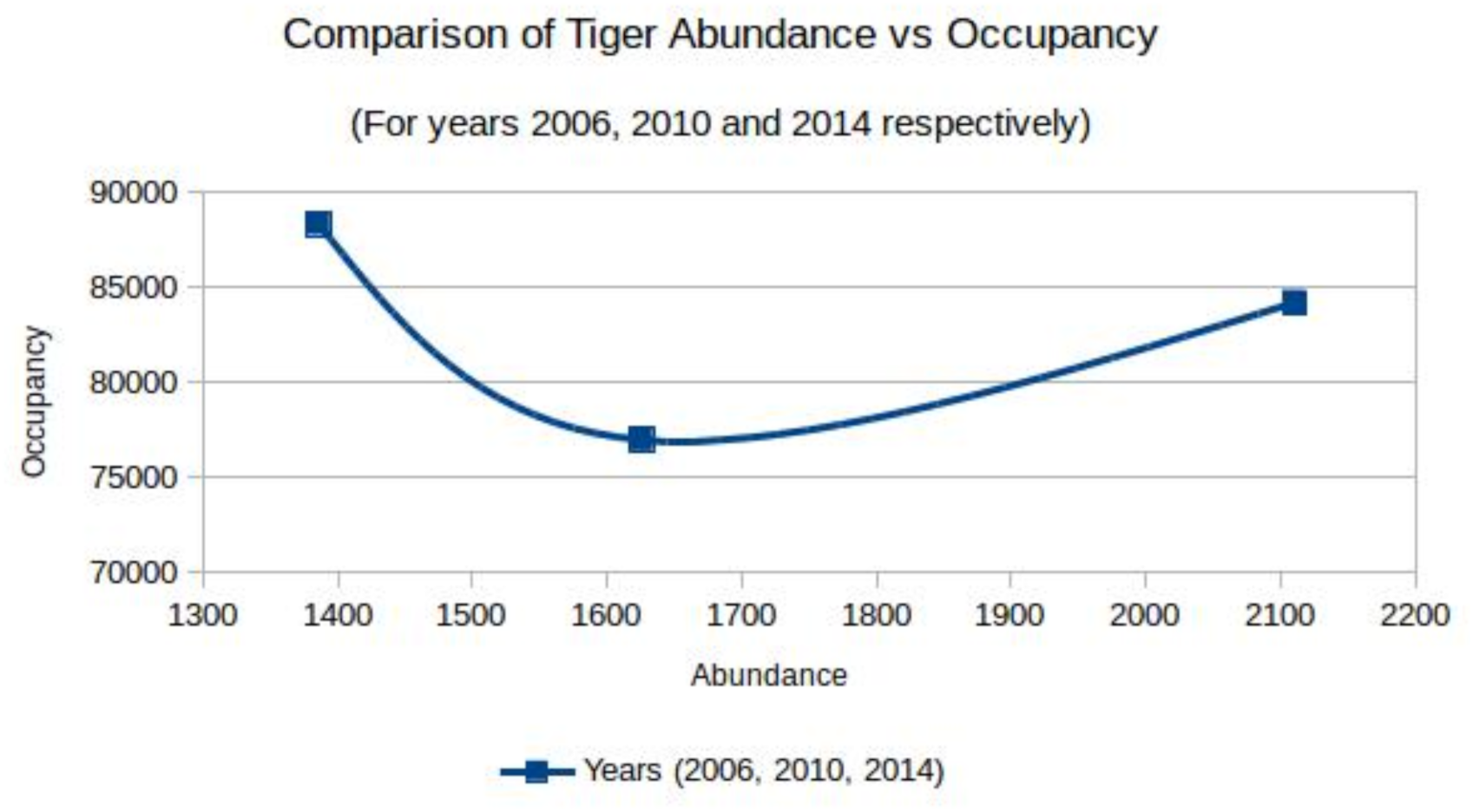
The occupancy-abundance relationship of India’s tigers. The concave upward relationship apparently challenges the existing monotonically-increasing, concave downward, occupancy-abundance relationship discussed in ecological literature.

Furthermore, long-term studies of tiger population dynamics using rigorous photographic capture-recapture surveys even in some better-protected tiger reserves of India (Karanth et al. 2006) and Thailand (Duangchantrasiri *et al.* 2016) demonstrate far lower annual rates of density increase (∼2-4%). If these survey outcomes are considered together the implication is that tiger populations in large, poorly-protected, low-prey density, sink landscapes exhibit higher growth rates than populations in better protected source populations (Karanth et al. 2016). Therefore, the results from the Indian tiger surveys stand in stark contrast to scientific understanding derived from the source-sink theory in population biology (Pulliam 1988), foundational to most global recovery plans for large carnivores including those for tigers in India (Walston 2010, NTCA 2012).

Recently, Harihar et al. (2017) analyzed these NTE survey results to show that increase of sampled areas in tiger ‘source sites’ among successive surveys led to decreases in tiger density. Hence, this study (Harihar et al. 2017) also contradicts implications from NTE surveys by showing that tiger population sinks are indeed performing worse than reported and supporting the proposals of Karanth et al. (2016) and Harihar et al. (2018), that tiger population recovery rates will be far slower than expected.

What are the reasons for these gross ecological anomalies that arise from Indian tiger surveys? The explanation by Darimont et al. (2018) is that ecological claims about political populations may often be disconnected from formal science. But the acceptance of the Darimont et al. (2018) explanation, without critical examination, may impede our gaining an understanding of tiger population biology from potential scientific serendipities (Wintle et al. 2010). Therefore, here we examine the basis of India’s claims on tiger numbers by assessing, in detail, the methods and models used to generate India’s tiger population estimates.

## DOUBLE-SAMPLING USING INDEX-CALIBRATION MODELS

The NTE survey method was developed and implemented in 2005 as India’s new official tiger monitoring approach (Jhala et al. 2008) after the failure of the previous ‘pugmark census’ method. Incidentally, the ‘pugmark census’ method persisted for three decades until it had to be suddenly abandoned only in 2005 due to its failure to detect the extirpation of an important tiger population (Sariska; Tiger Task Force 2005), followed by the extinction of another important tiger population (Panna; Special Investigation Team 2009).

The NTE survey method is based on the double-sampling experimental approach (Eberhardt and Simmons 1987). Double-sampling was developed because rigorous estimation of abundance at large spatial scales is often impractical because of ecological, environmental and logistical constraints (Eberhardt and Simmons 1987). When applied correctly, double-sampling involves the following steps:

(1) Random selection of a sample of sites from a larger pool of potential sites spread across a large region.

(2) Conduct of surveys at the sampled sites to estimate true animal abundance using a rigorous, reliable, method which is typically expensive, intensive and relatively difficult to implement (e.g photographic capture-recapture sampling, distance sampling).

(3) Conduct of a less rigorous, but practically feasible, field survey using an index of animal abundance (eg. number of animal tracks/km walked) at the selected sites as well as across the larger region.

(4) Development of an ‘index-calibration’ model (often a simple linear regression model) to establish a statistical relationship between true animal abundance (2) and its putative index (3).

(5) Estimation of animal abundance for the larger area using the index-calibration model (4).

Consequently, the reliability of results from the double-sampling approach will rest on the performance of the index-calibration model (step 4 from above) developed.

In the past, practical index-calibration experiments have yielded divergent results in terms of efficiency (see Gopalaswamy *et al.* 2015a,b and citations therein). Gopalaswamy et al. (2015a,b) mathematically modeled typical index-calibration models to analyze key contributors that produce such divergent outcomes in real world field surveys. In Appendix 1, without getting into all the mathematical complexities of Gopalaswamy et al. (2015a,b) we, heuristically, summarize the concepts of sampling-based overdispersion (SOD) and parameter covariation.

## INDEX-CALIBRATION MODELS OF INDIA’S TIGER SURVEYS

Based on mathematical and simulation-based explanations presented in Appendix 1, we reassess the inferences of empirical, large-scale, tiger surveys discussed in Gopalaswamy *et al.* (2015a,b). We elaborate on certain empirical details that were implicitly assumed in Gopalaswamy et al. (2015a,b) to contextualize the concepts for conservationists.

### (i) Assessing the predictive strengths of tiger sign index-calibration experiments

Gopalaswamy *et al.* (2015a,b) examined two different tiger sign index-calibration experiments (briefly labeled IC1 and IC2), which produced extremely divergent calibration successes. As a framework for statistical comparisons, they assumed that over India’s tiger occupied habitat of about ∼80000 km^2^ (Jhala et al. 2011a), there could be a potential pool of more than 400 sites each of ∼ 200 km^2^ size (approximately the size of sites used in the two experiments). In the two experiments, at each site, an estimate of tiger density was derived from photographic capture-recapture sampling (Karanth & Nichols 1998) from replicated surveys (see Karanth *et al.* 2004, Jhala *et al.* 2011b, for field work details).

At these sampled sites, tiger signs (scats and tracks) were counted by observers walking along trails to derive encounter rate indices (number of scats or track sets/km walked). These index count data, *S|N* were fitted to linear regression models by Ordinary Least Square (OLS) solutions. The first experiment (IC1), with a sample size of 21 sites, returned a high *R*^*2*^ estimate of 0.95 (as reported in Jhala et al. 2011a), whereas the second experiment (IC2), with a sample size of 8 sites, returned a low *R*^*2*^ estimate of 0.0004 (as computed using the lm function in R that uses the Eqn 4 from Kvalseth 1985 for estimating *R*^2^).

We note that the slope of these index-calibration relationships is *β=kp*\*, where *p*\* is the average detection probability per individual. Because the index based on tiger signs was computed from counts obtained in single sweep of each site, we set the value of *k=1.* Thus, making *β=p*\* in this case. If we apply the mathematical formula derived by Gopalaswamy *et al.* (2015a,b) for population *R*^*2*^ to the binomial model (less overdispersed case) we can obtain the estimate of detection probability *p*\*. This computed value is seen to be high for IC1 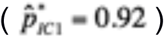 and low for IC2 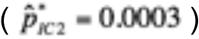. These two slopes are plotted as blue lines in Figure 4.

### (ii) Estimating the true value of p* for tiger sign index-calibration experiments

From the larger tiger distribution surveys conducted by Jhala *et al.* (2011a) and Karanth *et al.* (2011), the average *p*\* from these two surveys has an estimated value of 0.125 (represented by the red line in Figure 4), lying between the two blue lines.

This result clearly shows that sample sizes in both experiments (IC1=21 sites and IC2=8 sites) were far too small to accurately reflect the population characteristics. Secondly, the sampled sites selected non-randomly were not truly representative of the assumed larger pool of >400 sites because they both failed converge on the correct population estimate of average *p*\*. This implies that *both* these index-calibration models have poor predictive power across the wider spatial region of interest. Furthermore, in IC1, the 21 sites selectively excluded southwestern Indian region (see Jhala et al. 2011a,b). As seen earlier, the presence of large SOD makes index-calibration models very data-hungry, and any such selective, and potentially biased, sub-sampling of sites will compound the predictive inefficiency of these models.

### (iii) Factors likely to influence the potentially large variation in p*

We note that the tiger sign index-calibration models assume *k* is a constant and is equal to 1. In reality, *k* will be a function of the number of index values accumulated during the days prior to sampling and hence the constant *k* assumption may itself be unreal. For example, in drier forests (that cover about 50% of tiger habitats in India), tiger scats may remain intact for days prior to the counting, whereas they disappear rapidly in wetter regions. Although, Gopalaswamy *et al.* (2015a,b) did not derive explicit expressions for such variations in *k*, by assuming *k* is a constant, such an assumption will further, unrealistically, imply that the slope of *β* is entirely due to the variation in *p*\*. We also note that *p*\**=αp.* In this context, *p* refers to the detection probability of an individual tiger and its magnitude being determined primarily by the type of substrate (see Figure 3). Similarly, *α* represents the total fraction trails sampled. For example, in a photographic capture-recapture study, Karanth *et al.* (2004) demonstrated that detection probability *p*\* was much higher for tigers in the denser forests of Tadoba (0.174) and Bhadra (0.22), compared to open forests at Panna (0.039) or Bandipur (0.055) sites. This was possibly because the unknown proportion of trails used by tigers that were actually sampled, *α*, is relatively higher in denser forests with lower density of trails. We note here though that the value of *p* can potentially be very high only in exceptional circumstances, for example, detecting tiger tracks in snow in Russia (Miquelle et al. 2015) but *α* will continue to be dictated by the relative sampling effort per unit area.

**Figure 2.**
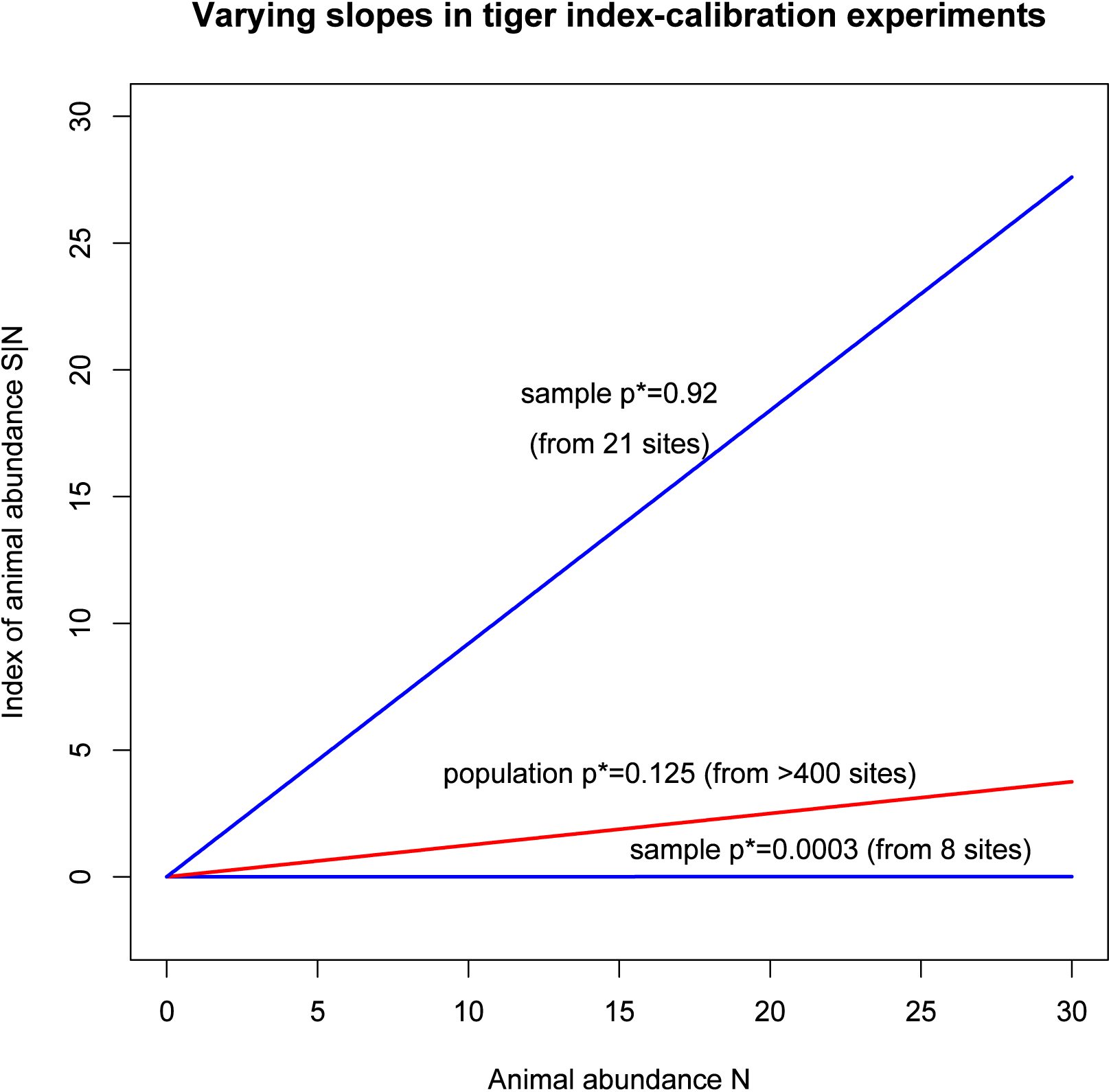
Illustration of the contrasting estimates of *p*\* under the binomial model of tiger index-calibration, *N* versus *S* (which is conditional on *N*). The lines are generated by the model *S|N ∼ Binomial(kN, p*\*), so that E(*S|N*)=*kNp*\*. The sampling occasion *k* is assumed to be a constant with a value of one. The two blue lines represent sample *p*\* estimates from two different tiger index-calibration experiments. The red line represents the line generated by a *p*\* estimate from an independent survey of the larger population of sites over Indian landscapes.

**Figure 3:**
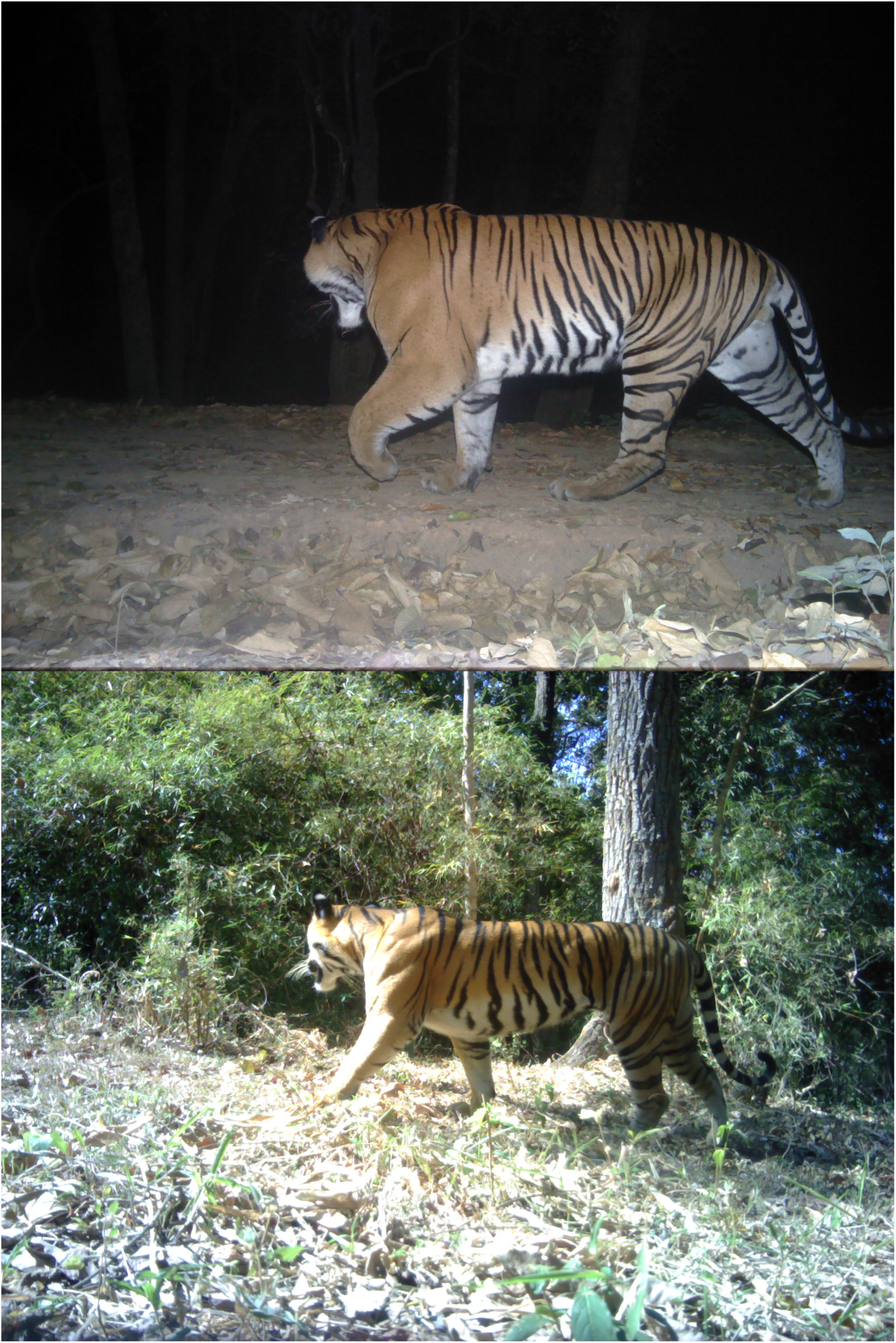
Photo-trapped images of tigers on contrasting substrate types. A dusty substrate (top) is conducive for detecting tiger tracks yielding a high detection probability *p*. In contrast, it is virtually impossible to detect tracks of tigers on leaf-littered, grassy, substrate types (bottom) yielding a low detection probability *p.* Picture courtesy of Ullas Karanth/WCS.

We also note that in both the above experiments IC1 and IC2, no particular spatial sampling design was employed to select trails. This factor also would additionally to contribute to causing biases in the estimate of *α*. Therefore, overall, the combined uncertainties of *k, α* and *p* are likely to contribute to the large variation seen in the value of *p*\*, resulting in the high overdispersion observed in such survey data.

## SAMPLING-BASED OVERDISPERSION IN INDIA’S TIGER SURVEY OF 2014

The resulting implication is that if SOD is not taken into account then any estimate of tiger abundance at the national scale will be non-robust and potentially flawed. To assess the generality of this conclusion, we evaluate estimates of tiger abundance from the previously unexamined NTE survey of 2014 (Jhala et al. 2015).

The calibration models developed during this survey comprised of a few environmental covariates in addition to tiger signs as explanatory variables to model tiger density. By the measure of *relative importance* of covariates (see Burnham and Anderson 2002), the survey results confirm that tiger sign index is the most important predictor of tiger density as this covariate is featured in the three main landscape models. This shouldn’t be surprising because signs of tigers are due to the presence of tigers. Whether treating tiger sign index as a relevant covariate in such modeling efforts is a separate question worthy of independent investigation. What is relevant to us here is whether there is SOD present in the relationship between tiger signs and tiger abundance.

In all the intensively monitored sites, the surveys estimated the *beta* coefficients corresponding to the tiger sign index covariate to be *betâ*_*SG*_ (*SÊ* (*betâ*_*SG*_)) = 0.1(0.06), *betâ*_*CIEG*_ (*SÊ* (*betâ*_*CIEG*_)) = 0.258(0.028) and *betâ*_*WG*_ (*SÊ* (*betâ*_*WG*_)) = 1.01(0.08), where SG, CIEG and WG correspond to abbreviated forms of Shivalik-Gangetic Plains, Central-Indian and Eastern Ghats and Western Ghats, respectively. We note here that the definition of *beta* in these reports will differ from our definition of β earlier in that *beta* is meant to represent the rate of change of animal density for a unit increase in the signs detected. However, our purpose is to investigate SOD and parameter covariation and these estimates of *beta* serve that purpose well enough.

The full mathematical specification of the model used in Jhala et al. (2015) is not available. However, from the model coefficients reported, they appear to be generated using the default log-linear model in the package secr (Efford 2019). Therefore, the above *beta* estimates must be back-transformed exponentially for appropriate interpretation. Accordingly, one unit increase in the tiger sign index results in a corresponding *exponential* increase in tiger density of (i) 10.5% in the Shivalik-Gangetic Plains (ii) 29.4% in the Central Indian and Eastern Ghats landscape and (iii) a massive 174.6% increase in tiger density in the Western Ghats. Such a massive variation in the influence of tiger sign index on tiger density demonstrates the presence of enormous SOD inherent in the population. Interestingly, these estimates of *beta* indicate that the non-linear nature of the relationship between tiger sign index and tiger density is very pronounced, perhaps indicating a strong interaction between *p* and/or *α* and *N* as discussed earlier.

Such variations arising from SOD mean that when these *beta* estimates are utilized to estimate tiger abundance over wider regions (eg: at regional and national levels) or used to assess changes in tiger abundance over time, the resulting trends can essentially lack any real ecological meaning. And the large variation in the extent of non-linearity further weakens predictions of tiger abundance at regional and national levels.

## CONSERVATION IMPLICATIONS

The *beta* estimates reported in Jhala et al. (2015) *strongly confirm* the presence of a high degree of SOD and parameter covariance in tiger index-calibration experiments used in Indian tiger surveys. The temporal variations in the *beta* estimates and the unpredictable changes in the form of the index-calibration relationship itself make the prediction of animal abundance at large spatial scales very unreliable. We conclude that changes in the tiger population size and occupancy reported from Indian tiger surveys, which are so anomalous in the context of ecological rationale (see *Introduction*), are outcomes from an unreliable method used which spuriously seems to challenge existing understanding of animal population dynamics in ecology.

There are several management implications that arise from our analyses. It is essential that the raw data from the past NTE surveys (Jhala et al. 2008, Jhala et al. 2011a, Jhala et al. 2015) be thoroughly re-analyzed to account for the hitherto ignored underlying SOD and parameter covariance we have we have uncovered here. Only such a re-analysis can correct these tiger population estimates, by fully recognizing the true underlying uncertainties that have been previously ignored. We note that, analytically, the SOD problem can at least be reduced partially by accounting for spatial random effects (Dey et al. 2017). More importantly, the understanding of the effects of SOD provides an opportunity for conservation agencies to introspect deeply about properly designing monitoring programs keeping in view suggestions of Nichols and Williams (2006) given the enormous resources (manpower, time and money) spent on Indian tiger surveys as detailed in Jhala et al. (2015).

## CONSERVATION OUTCOMES: DOES SCIENCE INFLUENCE POLICY?

Accepting the premise that species conservation programs should be based on science and evidence, the conservation implications discussed above can serve as a template for future tiger monitoring policies. But, as discussed in Darimont et al. (2018), this often will not be the case with political populations.

Our understanding of the presence of large SOD in India’s official tiger estimation approach was based on the 2010 survey results and presented in Gopalaswamy et al. (2015a). Coincidentally, this publication appeared only a month after India announced that its tiger population had risen by 30% during the years 2010-2014.

Surprisingly, instead of seeking more information, studying implications of Gopalaswamy et al. (2015a) in detail, or by engaging in a formal scientific debate, some scientists and officials associated with the NTE survey, rushed to the journal demanding, summarily, the retraction of Gopalaswamy et al. (2015a) (Vishnoi 2015, Kempf 2016). Given that our analysis of the NTE survey of 2014 discussed here further *strengthens* arguments advanced by Gopalaswamy et al. (2015a,b), this conservation outcome of demanding retraction is perplexing.

We note with some concern that in spite of the presence of a body of scientific evidences pointing towards major concerns about India’s tiger population rise claims (Gopalaswamy et al. 2015a,b, Karanth et al. 2016, Harihar et al. 2017, Harihar et al. 2018) major international conservation agencies, such as the Global Tiger Forum (GTF), Global Tiger Initiative (GTI), and the World Wide Fund for Nature (WWF), continued to endorse claims of success made by Indian Tiger Surveys (WWF 2016). Consequently, India’s tiger conservation budget jumped from USD $70 million to $144 million in 2016 to reward this achievement (PTI 2016).

Of even greater concern is the fact that index-based monitoring methods continue to be uncritically employed by India’s National Tiger Conservation Authority and Wildlife Institute of India (Jhala et al. 2017) and even being further promoted by GTF and by the GTI in other tiger range countries that are just initiating national monitoring programs (see Dey et al. 2015 for an example of from Bangladesh). Similarly, Nepal also claimed that their tiger numbers doubled in a relatively short period of time (Davis 2018) without adequate scientific support.

More recently, Qureshi et al. (2018) archived in a public preprint repository a critique of Gopalaswamy et al. (2015a,b), but at the same time supporting claims of the NTE survey of 2014 (Jhala et al. 2015). Since we demonstrate that Jhala et al. (2015) only buttresses the scientific findings of Gopalaswamy et al. (2015a,b), especially with regards to SOD and parameter covariance, we find that this critique (Qureshi et al. 2018) too appears to be a paradoxical conservation outcome (Gopalaswamy 2019), and contrary to science-based implications.

## DISCUSSION

As we elucidate in this article, the combined phenomena of sampling-based overdispersion and parameter covariance can induce a large amounts of uncertainty in predictions of animal abundance over large spatial and temporal scales. This implies that the claims of a 58% tiger population rise in India over the past 8 years (from 2006-2014) based of estimates from the three NTE surveys (Jhala et al. 2008, Jhala et al. 2011a, Jhala et al. 2015) lack reliable scientific support.

Our analysis of these additional sources of uncertainty, however, helps in explaining the ecological paradoxes (see *Introduction*) that the survey results lead to. For example, the unusual concave upwards relationship between tiger occupancy and abundance (Figure 1; derived from the summary table in Jhala et al. 2015), the reversal of the source-sink mechanism (Karanth et al. 2016, Harihar et al. 2017) and the unusually the high estimates of tiger population growth rates in source populations (Karanth et al 2006, Duangchantrasiri et al. 2016) all appear to be consequences of inferential problems arising from sampling-based overdispersion and parameter covariance.

In general, we believe that such misreading of population growth patterns resulting from inferences based on vague survey methodologies can be detrimental to wildlife conservation. Just over a decade ago a similar disregard to scientific findings, using an earlier flawed survey methodology known as the pugmark census (Karanth et al. 2003) had hidden real tiger population collapses in India. At that time, India’s tiger numbers were reported to have reached 3642 individuals (Ramesh 2008). This was followed by abandonment of the census method, as tiger populations collapsed from a wave of poaching in two key reserves in India (Tiger Task Force 2005, Chundawat 2018).

The current claim of an upward trend in tiger numbers is reminiscent of that previous conservation ‘bubble’. Based on our results and more general suggestions of (Nichols and Williams 2006), we question the very purpose of conducting such massive, resource intensive surveys across large regions without first addressing challenge of SOD and parameter covariance, which remains challenging if not intractable.

We note that at the high profile Global Tiger Summit in St. Petersburg in 2010, *doubling* the global number of wild tigers by 2022 was proclaimed as the goal and financial commitments of about US $ 330 million were pledged (Watts 2010). We worry that large financial commitments in species conservation initiatives may subconsciously create social pressures on conservation agencies leading to motivation or cognitive bias (Kunda 1990, Kahneman 2013) influencing either the survey design itself or the reporting of results from surveys. For example, by claiming rising tiger numbers, Indian conservation agencies obtained an immediate increase in their tiger conservation budgets (PTI 2016). And when non-robust monitoring survey methodologies (eg: with inherently large SOD and/or parameter covariance) are employed, they can even dangerously legitimize any popular claim about population trends because such estimates can understate the true, but much wider, confidence intervals. We observe the similarity of this situation to the 4-way categorization by Pielke (2007) corresponding to the category of “high scientific uncertainty, popular choice”.

We therefore stress the importance of structuring a sound monitoring program in species conservation initiatives (Nichols and Williams 2006, Karanth and Nichols 2017). While claims about population increases or decreases may meet the goal of increasing public support and assist in raising more funds for conservation, we argue that this line of reasoning can be hugely detrimental to species conservation. First, such an approach will tend to benefit the most *advertised* conservation strategy as opposed to the most *effective* one. Consequently, those invested in solving on-ground conservation or scientific problems could potentially be pressured into investing time and effort in marketing and outreach. Second, monitoring programs that do not truly advance scientific knowledge will undermine the entire discipline of field ecology itself. For example, with respect to our above example with tigers, the unexplainable concave upward relationship of tiger occupancy-abundance dynamics (Figure 1) or the inexplicable accelerated growth rate of tiger populations only raises more unsolvable ecological questions rather than providing good answers.

We argue that when conservationists fail to keep pace with novel scientific methodologies, any claimed estimate will prima facie be non-robust. If massive changes in tiger numbers is attributed to change in methods (eg: drop in India’s tiger numbers from 3642 tigers (Ramesh 2008) to 1411 tigers in 2006 (Jhala et al. 2008)) or due to unreliable methods (as analyzed here) it has little value to either science or conservation because monitoring can no longer provide useful information to conservationists in real time.

Our article attempts to convey and contextualize the mathematical and empirical findings of Gopalaswamy et al. (2015a,b) to conservationists, especially in the context of India’s claims of tiger population rise. But there is vast scope for further research on this theme. If we ignore the presence of SOD and parameter covariation for the moment, it is ecologically interesting to assess these tiger population dynamics in the context of biological overdispersion (May 1978). Since Jhala et al. (2015) and Gopalaswamy et al. (2015a,b) demonstrate the difficulty of applying double-sampling in practice for tigers, it becomes relevant to derive the parameters for the population level overdispersion caused by variation in detection rates, even though the relevance of using indices for regional population level estimation may remain futile (see Belant et al. 2019, in the case of lions in Serengeti).

With respect to tigers, we recommend that monitoring investments be targeted to reliably understand the drivers influencing vital rates (survival, recruitment and movement) at critical tiger source populations (Karanth et al. 2006, Duangchantrasiri et al. 2016, Walston et al. 2010). Complementarily, we recommend landscape scale, sign surveys (conducted once in 4-5 years) to track and understand the mechanisms of tiger range contractions, expansions and connectivity (Karanth and Nichols 2017).

More generally, we propose that animal monitoring programs, particularly of political populations such as tigers, be designed and implemented *entirely* to answer sound scientific and management objectives (Krebs 1991, Nichols and Williams 2006) rather than be influenced by social or fund raising pressures from extraneous sources. Such a focus will help prevent charismatic large carnivore populations to avoid the risk of being stigmatized as a political population (Darimont et al. 2018).

## Ackowledgements

We thank the Nigel Yoccoz, two anonymous reviewers and the Associate Editor for helpful suggestions on the earlier version of this paper. AMG thanks the Indian Statistical Institute, Bangalore Centre and Wildlife Conservation Society, New York for funding support. We thank David W. Macdonald and Tim Coulson for encouraging us and providing administrative support to AMG during his time at Oxford. We thank Bob May, Jim Nichols, Sari C. Cunningham, Devcharan Jathanna and Varsha S. Shastry for helpful comments and assistance with the drafts.

## SUPPLEMENTAL INFORMATION

## APPENDIX 1) INDEX CALIBRATION MODELS: Summary of concepts from Gopalaswamy et al. (2015a,b)

### Basic Statistical Concepts

#### The binomial and beta-binomial index-calibration models

As in Gopalaswamy et al. (2015a,b), we define N as the true animal abundance at a site and S as the corresponding index of abundance measured at the site. We assume N is known noting that Gopalaswamy et al. (2015a,b) did not require this simplifying assumption. We assume that a selected site is sampled over k occasions independently. If p* is the probability of detecting an individual animal at this site during one occasion, then the index over k occasions is modeled as S|N,k,p* ∼ Binomial(kN, p*). S|N,k,p* (or S|N, for simplicity) is read as S conditional on N, k and p*. This binomial model has the expected value (or expectation) E(S|N)=kNp*. For example, if we conduct a survey of animals at a single site on 3 occasions, with the true abundance being 20 animals, and the detection probability per occasion being 0.1, then the average count from such a survey is kNp*=(3)(20)(0.1), or 6 encounters. For a large number of repeated experiments of this kind, the experiment-to-experiment variation in counts at this site (with N=20) is described by Var(S|N)=kNp*(1-p*)=6(1-0.1)=5.4.

In the above example, we have assumed detection probability p* to be unvarying. In reality, there are many sources that can induce variation in p* either temporarily or spatially. To account for such variation, Gopalaswamy et al. (2015a,b) developed a beta-binomial version of the model, so that S|N,k,a,b ∼ Beta-binomial(kN, a, b). Here, the detection probability is described as a random quantity drawn from a beta distribution with shape parameters a and b. Under such a model, the variation in the animal counts obtained will always be larger than in the binomial case (implying that Var(S|N) > 5.4 if there is variation in p*). In either case, the variance of the count increases when N increases. This critical phenomenon is defined statistically as overdispersion. We specifically define this phenomenon as ‘sampling-based overdispersion (SOD)’ to distinguish it from ‘biological overdispersion’ traditionally defined in ecology to represent the heterogeneity in abundances over space (May 1978).

#### (ii) Slopes of the index-calibration experiments

A good index-calibration experiment involves conduct of similar surveys at multiple sample sites, with different values of N. These counts result in the index, S|N, from all these sites. For a fixed value of k and mean p*, the average slope, say β, of this relationship is given by kp* for the binomial model, and k[a/(a+b)] for the beta-binomial model. With parameters defined earlier, for example applying a coefficient of variation in p* of 0.4 for the beta-binomial case, we can simulate multiple data points for two imaginary index-calibration experiments (see Figures 1a,1b). The divergent, flash-light like, spread of these data points (in blue circles) indicates the extent of SOD inherent in such data.

#### (iii) Measuring predictive strength in index-calibration experiments

The strength of an index-calibration experiment can be assessed either from graphical visual assessments or using formal statistical measures. Gopalaswamy et al. (2015a,b) used the coefficient of determination R^2^ (a goodness-of-fit measure commonly used in the applied sciences) to assess predictive strengths index calibrations under both the binomial model and beta-binomial models. They showed that strong predictive relationships (R^2^ is close to 1) are obtained when detection probability p* is high as well as unvarying. In our simulated experiments, the estimated R^2^ is lower for the beta-binomial model (0.46) model compared to the binomial (0.63) model, because of its greater SOD. Further, Figure 1c shows how the R^2^ measure increases from 0.46 to 0.54, when the slope of the relationship, determined by average detection probability p*, increases from 0.1 to 0.2 in the beta-binomial model.

### Field sampling concepts

#### Random and representative selection of sites for double-sampling

The basic statistical concepts and mathematical formulae applied by Gopalaswamy et al. (2015a,b) assume that the sample size (number of index-calibration data points) is infinite or very large. In practice, this is unrealistic because only a few data points are usually selected for fitting the index-calibration model. But it is essential that the selection of data points should at least representatively retain the same slope and ‘flash-light’ like SOD shown in Figures 1a, 1b and 1c. Only with such random site selection can an investigator ensure that model predictions from sampled points can be extended to generate abundance estimates to larger spatial scales using the double-sampling approach. Therefore, the assumption that the sites sampled do representatively capture the overdispersion pattern in this manner becomes critical to extend inferences about animal abundance to wider areas.

#### (ii) Deconstructing the detection probability p*

The detection probability p* defined here is actually the product of two probabilities: α and p, so that p*=αp, where α is the proportion of the area within each site that is actually sampled, and p is the probability that an animal within the sampled area in that site is detected during the survey (Williams et al. (2002)). Hence, variation in either α or p will inevitably induce variation in p* (Elliot and Gopalaswamy (2017), Karanth and Nichols (2017)).

### (c) Data analytic concepts

#### Fitting index-calibration models using overdispersed data

Ideally these index-calibration models confronting overdispersed data should be defined by likelihood functions for specific data-generating cases (binomial or beta-binomial) (see Richards 2008). However, animal monitoring studies often apply standard linear regression models using ordinary least square (OLS) solutions prescribed in standard textbooks (e.g. Sutherland 2006). There is an inherent problem in doing this, as we illustrate in Figure 3. For p*=0.5, the expected value of S|N is indicated by a solid dark green line. Let us assume that the average p* comes from an underlying beta distribution, with a coefficient of variation = 0.4. In sign-based indices, k may be unknown and there is an inherent identifiability issue between k and p*. So, we assume that the number of sampling occasions k is fixed and set to 1 (see Gopalaswamy et al. (2015a,b) for the limiting case when k =∞). The circles, distributed around this line, represent one simulated outcome from a set of 20 imaginary data points. When a standard linear regression analysis is conducted on data to relate the variables S|N and N, an imaginary straight line is constructed (the solid orange line) through the data points. Visually, the placement of this line involves finding an alignment, which will minimize the least distance between each data point and the line. This is called the OLS solution (Casella and Berger 1990). We can use this fitted line to draw inferences about the regression parameters (the slope and intercept) of the linear relationship between S|N and N. We conducted such a standard linear regression analysis by OLS on our simulated set of data points and plotted the regression line (in solid orange) along with its associated confidence intervals (in dashed orange).

We notice here (Figure 2) that while the index S tracks the variation in N reasonably well, the absolute variation in S|N itself is quite large, especially when N increases. For this particular simulation, the OLS solution leads to an overestimation of the slope by a substantial degree (45.4%). If this fitted regression line were to be used for making predictions of animal abundance over large scales, seriously biased estimates would result. What is worse is that the direction and magnitude of the bias depends on the true value of N. Further, we note that the estimated 95% confidence intervals do not even bind the true expected value (dark green line) in many regions of the graph. Where does this inconsistency come from? The answer is found in statistical theory (Kruskal 1968). It turns out that OLS fits are inefficient in the presence of overdispersion – in our simulated case due to the presence and variation in p*. In fact, this key problem is well recognized in the econometrics literature (Hayashi 2000). This mathematical result means that if overdispersion is present (as it indeed is in most animal survey data sets), abundance predictions employing double sampling approaches using standard linear regression analysis by OLS will not reflect the true underlying uncertainty (eg: poor coverage probability).

#### (ii) How parameter covariance induces artificial non-linearity in index-calibration relationships

There is an additional problem affecting the index-calibration method. Often, an inherent covariance can exist between model parameters. The binomial model is denoted as: E(S|N)=kNαp=(kαp)N=βN. Here, we notice that there are three sampling-related parameters, k, α and p, and one ecological parameter, N. In practice, if there are underlying ecological or sampling relationships among some of these parameters, the index-calibration relationship will take a non-linear form. In such cases, linear regression models are no longer applicable and their predictions from index calibration data will further be at fault.

## Figure Legends

**Figure 1.**
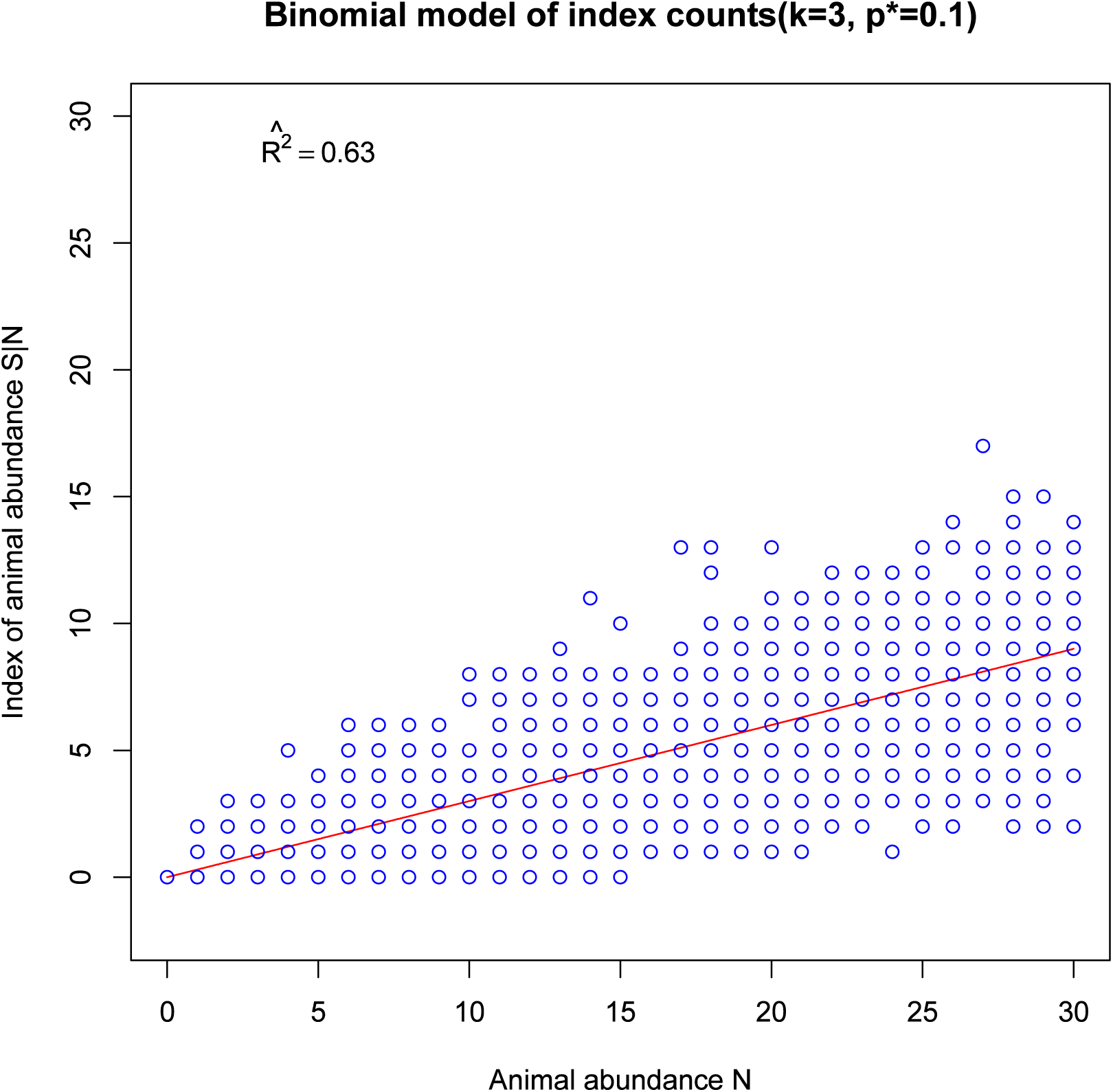
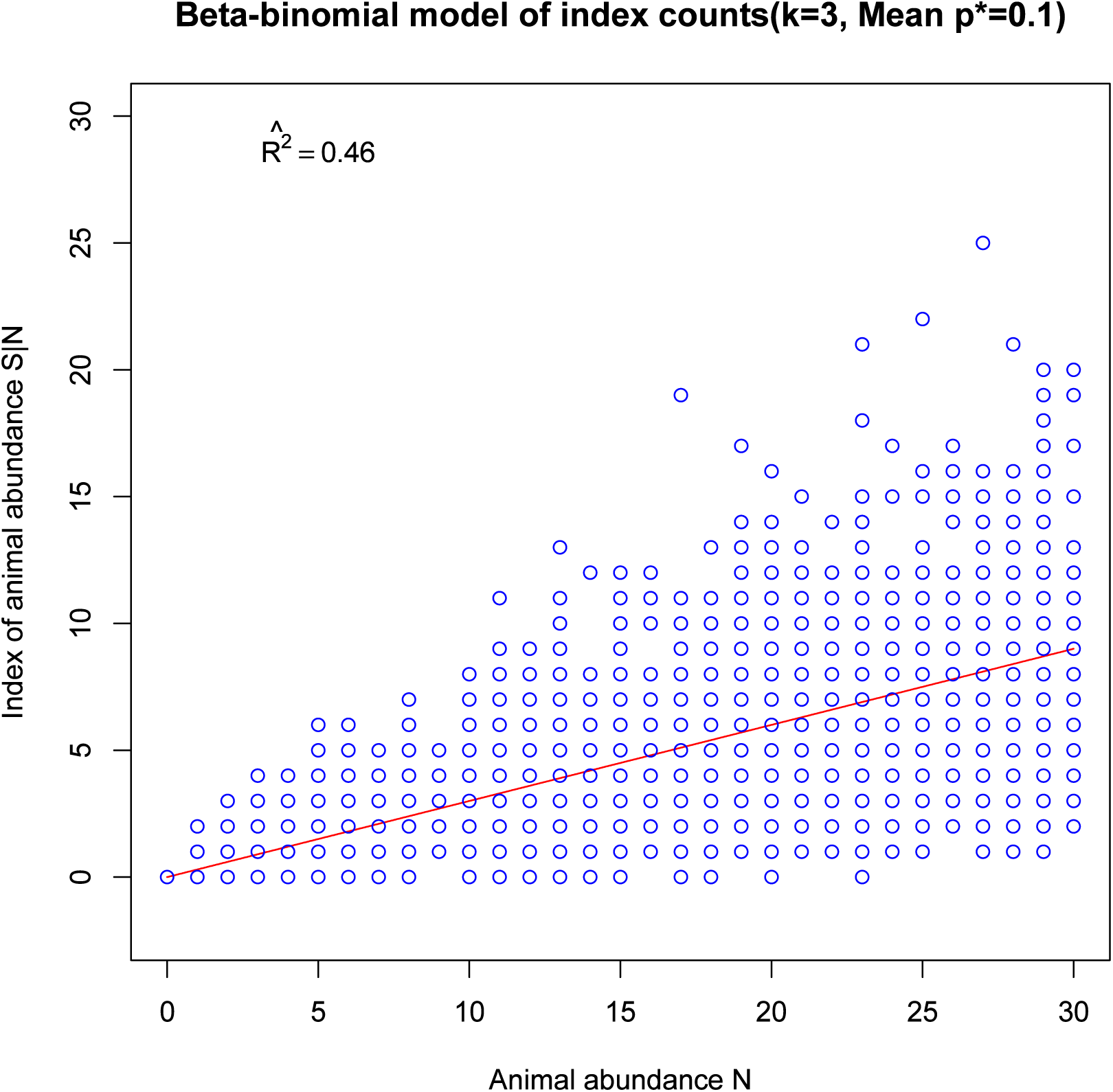
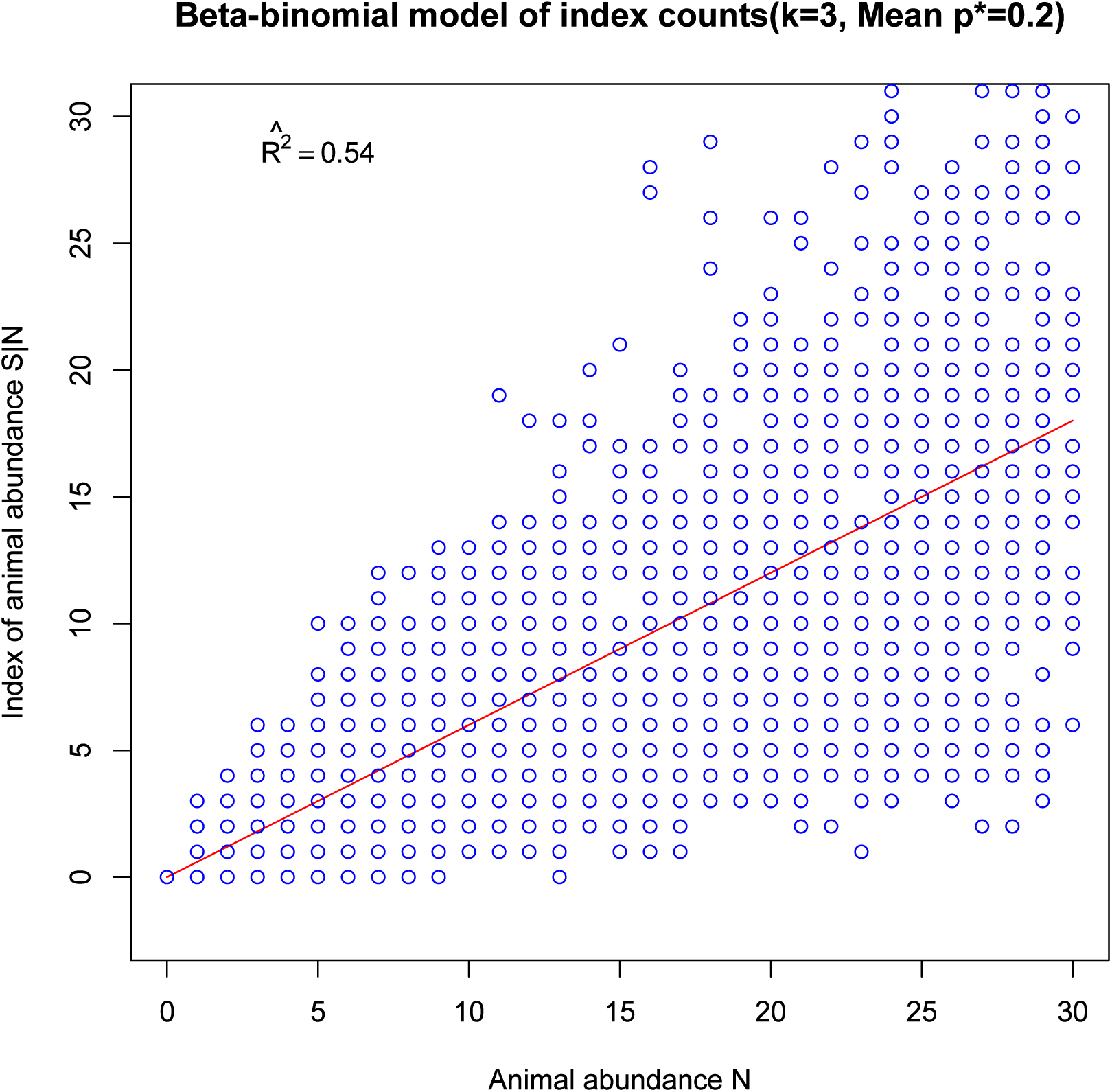
Simulation results for hypothetical index-calibration experiments (abundance index *S|N* versus *N*) with large sample sizes (2000 data points) are depicted here for the (a) binomial model and (b & c) beta-binomial models. The number of sampling occasions is fixed at *k=3.* We show that the estimated *R*^*2*^ value drops from (a) to (b) when a coefficient of variation of 0.4 is applied to the average detection probability parameter *p*\* (set at 0.1 for both cases). Similarly, the *R*^*2*^ value increases from (b) to (c) when the average detection probability parameter *p*\* increases from 0.1 to 0.2. The red lines depict the expected index-calibration relationship, E(*S|N*)=*kNp*\* for the binomial model and E(*S|N*) = *kN*[*a/*(*a+b*)] for the beta-binomial model. The parameters *a* and *b* are estimated from the given coefficient of variation by the formulae provided in Gopalaswamy *et al.* (2015a,b).

**Figure 2.**
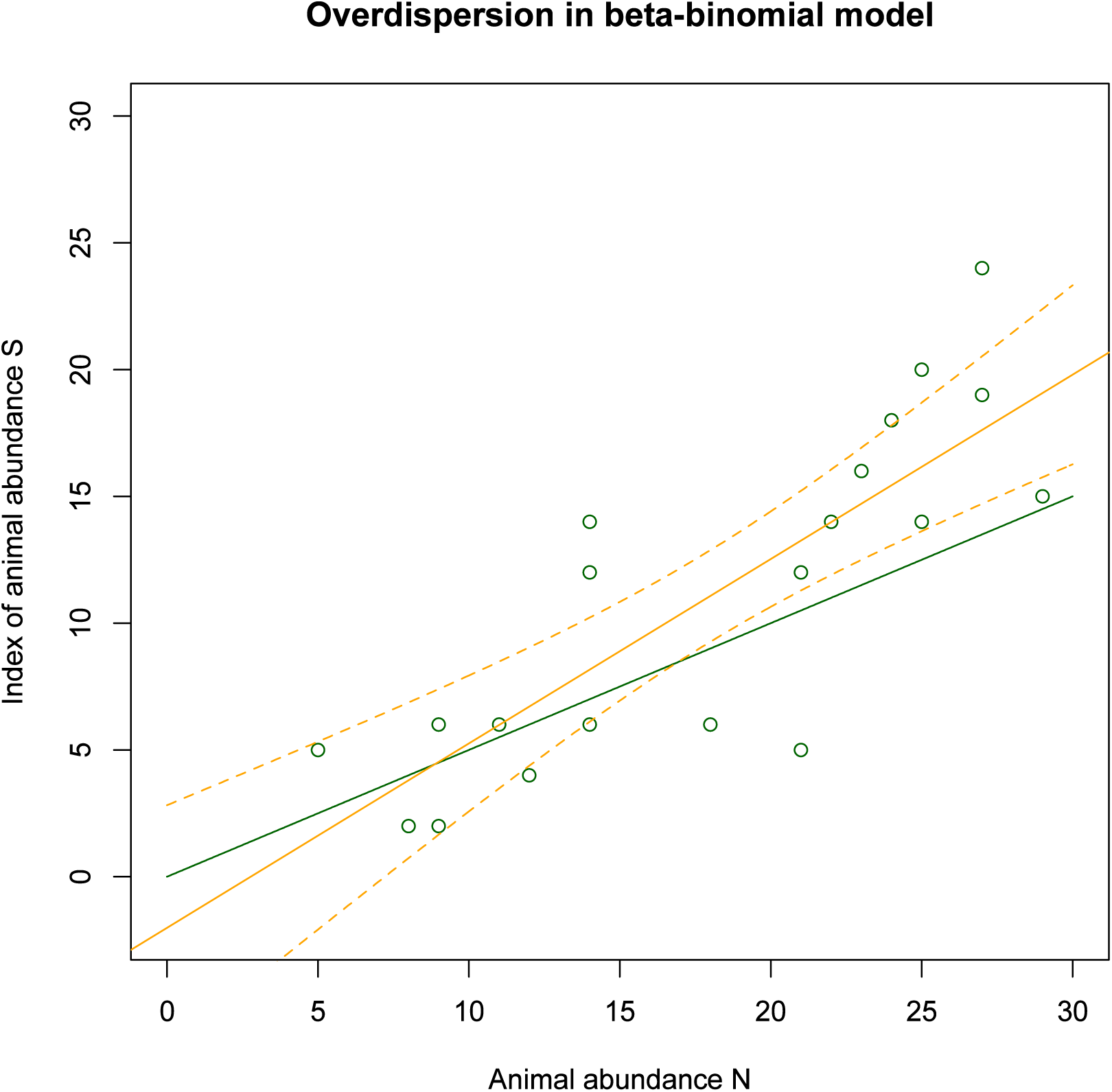
Illustration of the extent of dispersion under the beta-binomial model of index-calibration, *N* versus *S* (which is conditional on *N*). The overdispersion is caused by the presence of a variable detection probability, so that *S|N ∼ Beta-binomial(kN, a, b)* with the corresponding *a* and *b* set to reflect a coefficient of variation of 0.4 around the average *p*\*. The sampling occasion *k* is assumed to be a constant with a value of one. The dark green circles represent a random selection of data points from the specified distribution, where mean *= Na/(a+b)* and variance =*[Nab(a+b+N)]/[(a+b)2(a+b+1)]*, for average p*=0.5.

